# Centriole-less pericentriolar material serves as a microtubule organizing center at the base of *C. elegans* sensory cilia

**DOI:** 10.1101/2020.08.20.260414

**Authors:** Jérémy Magescas, Sani Eskinazi, Michael V. Tran, Jessica L. Feldman

## Abstract

During mitosis in animal cells, the centrosome acts as a microtubule organizing center (MTOC) to assemble the mitotic spindle. MTOC function at the centrosome is driven by proteins within the pericentriolar material (PCM), however the molecular complexity of the PCM makes it difficult to differentiate the proteins required for MTOC activity from other centrosomal functions. We used the natural spatial separation of PCM proteins during mitotic exit to identify a minimal module of proteins required for centrosomal MTOC function in *C. elegans*. Using tissue specific degradation, we show that SPD-5, the functional homolog of CDK5RAP2, is essential for embryonic mitosis while SPD-2/CEP192 and PCMD-1, which are essential in the zygote, are dispensable. Surprisingly, although the centriole is known to be degraded in the ciliated sensory neurons in *C. elegans* [1-3], we find evidence for “centriole-less PCM” at the base of cilia and use this structure as a minimal testbed to dissect centrosomal MTOC function. Super-resolution imaging revealed that this PCM inserts inside the lumen of the ciliary axoneme and directly nucleates the assembly of dendritic microtubules towards the cell body. Tissue-specific degradation in ciliated sensory neurons revealed a role for SPD-5 and the conserved microtubule nucleator γ-TuRC, but not SPD-2 or PCMD-1, in MTOC function at centriole-less PCM. This MTOC function was in the absence of regulation by mitotic kinases, highlighting the intrinsic ability of these proteins to drive microtubule growth and organization and further supporting a model that SPD-5 is the primary driver of MTOC function at the PCM.

## Results

Microtubules perform numerous cellular functions, including chromosome segregation and intracellular transport, that require spatial organization imparted by cellular sites called microtubule organizing centers (MTOCs). The best-studied MTOC is the centrosome, a non-membrane bound organelle composed of two barrel-shaped microtubule-based centrioles surrounded by a layered network of pericentriolar material (PCM; [4-6]). The MTOC activity of the centrosome cycles with the cell cycle, peaking during mitosis when the PCM expands due to regulation by mitotic kinases and phosphatases to recruit microtubules.

The molecular complexity of the PCM and its regulation has obscured the identity of proteins that function in microtubule organization *per se* rather than other aspects of centrosome biology. In *C. elegans*, the PCM is relatively simple and composed of two main scaffolding proteins, SPD-2/CEP192, SPD-5 (the functional homologue of CDK5RAP2), and a recently discovered coiled-coil protein PCMD-1 [7]. These proteins have largely been characterized in the zygote where they are interdependently recruited to the centrosome and, together with AIR-1/Aurora-A, localize the conserved microtubule nucleation complex γ-TuRC [8-14]. Despite the simplicity of *C. elegans* PCM, the direct intramolecular interactions that ultimately build and anchor microtubules are largely unknown. We therefore aimed to understand which PCM proteins impart the MTOC function of the centrosome in multiple cellular contexts.

Our recent analysis in early dividing embryonic cells found evidence for distinct subcomplexes within the *C. elegans* PCM [5], only some of which are spatially associated with MTOC function. In particular, we found that the PCM is organized with an ‘inner’ sphere that localizes all PCM proteins and an ‘outer’ sphere that colocalizes SPD-5 and proteins like γ-TuRC required for microtubule growth and assembly. These subcomplexes are not only spatially separated but also appear to be functionally distinct as MTOC function segregates with outer sphere proteins during PCM disassembly in mitotic exit: proteins in the outer sphere rupture into “packets” which co-localize with dynamic microtubules. Thus, SPD-5 is the central PCM component in the outer sphere and in packets (Figure 1A and1B; [5]). Conversely, SPD-2 and the recently identified PCMD-1 protein are spatially restricted closer to the centrioles (Figure 1A and S1), and do not localize in packets during disassembly (Figure 1B; [5]). 3-Dimensional Structured Illumination Microscopy (3D-SIM) imaging of PCM revealed that SPD-2 was concentrated more centrally (Figure 1C) while SPD-5 formed a broad perforated matrix. During packet formation, SPD-2 localization was restricted to two toroids that we assume surround the centrioles as the localization resembled that of the ‘naked’ centrioles in the *C. elegans* gonad. In addition to localizing to packets, SPD-5 also localized as a toroid, however, the SPD-2 and SPD-5 toroids appeared to be offset and stacked (Figure 1C and 1D).

**Figure 1.**
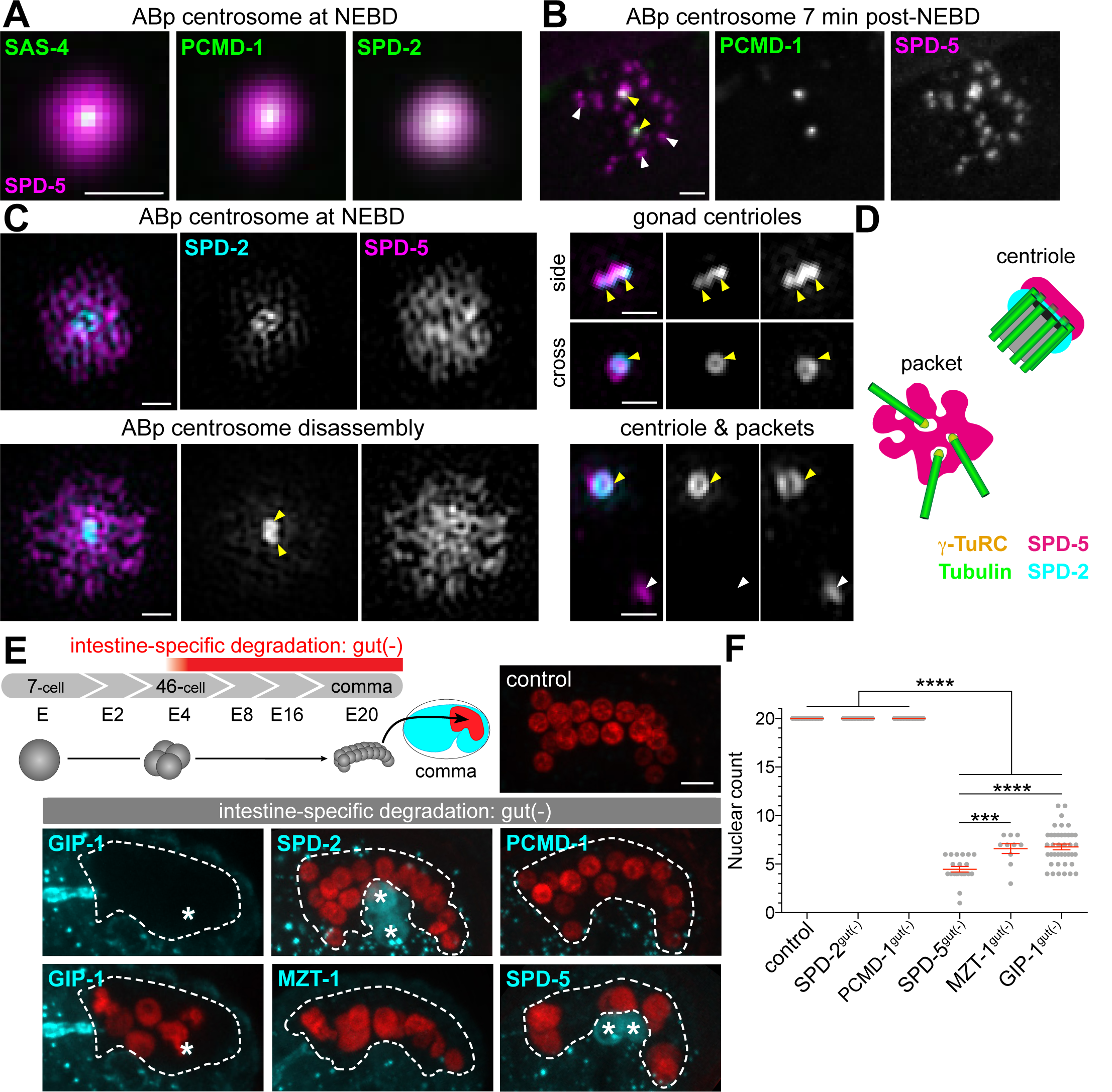
MTOC function at the centrosome is driven by SPD-5 dependent subcomplexes independent of SPD-2 or PCMD-1, see also Figure S1. A-B) Live confocal imaging of indicated proteins in the ABp centrosome from a 4-cell embryo at nuclear envelope breakdown (NEBD) or 7-minutes post-NEBD. Scale bar= 5µm. B) PCMD-1 does not localize to SPD-5 positive packets (white arrowheads), only to separated centrioles (yellow arrowheads). C) 3D-SIM imaging of centrosome structures at indicated stages. Centrioles (yellow arrowheads) and packets (white arrowheads) are indicated. Scale bar= 500nm. D) Cartoon representing the separation of function of PCM proteins, with MTOC function specifically associated with packets. E) Top: Cartoon depicting the development of the *C. elegans* embryonic intestine (left) relative to the E4-stage onset of *elt-2p* promoter expression. Embryonic intestines will achieve 20 cells by comma stage, which can be visualized using an intestine-specific histone::mCherry. Below: Intestine-specific ZIF-1 expression (*elt-2p::zif-1*) effectively degrades indicated ZF:GFP tagged proteins (cyan). * indicates germline cells or non-intestinal signal. Scale bar= 5µm. F) Analysis of intestinal nuclear number at comma stage: control: mean= 20±0.0 S.D. nuclei, n=20; SPD-2: mean= 20.0±0.0 S.D. nuclei, n=20; PCMD-1: mean= 20±0.0 S.D. nuclei, n = 20; SPD-5: mean= 4.5±1.3 S.D. nuclei, n= 21; MZT-1: mean= 6.6±1.6 S.D. nuclei, n= 10; GIP-1: mean= 6.8±1.9 S.D. nuclei, n= 40. Tukey’s multiple comparisons test, *** p-value=0.0008, **** p-value<0.0001.

### Embryonic mitosis relies upon SPD-5 but not SPD-2 or PCMD-1

Based on our localization studies, we hypothesized that MTOC function at the centrosome is controlled by a SPD-5-dependent protein module that functions independently of SPD-2 and PCMD-1. To test this hypothesis, we depleted essential centrosome components in late embryonic cell divisions using a tissue-specific degradation system to deplete endogenous SPD-5, SPD-2, PCMD-1 and γ-TuRC components GIP-1/GCP3 and MZT-1/Mozart1 from the 4-cell stage embryonic intestine (Figure 1E, ‘gut(-)’, ‘E4’; [15-17]). Intestinal cells all derive from the E-blastomere through successive rounds of divisions leading to an invariant 20-cell intestine in control embryos (Figure 1E and 1F). γ-TuRC depletion resulted in significantly fewer intestinal cell nuclei (Figure 1E and 1F, GIP-1^gut(-)^ and MZT-1^gut(-)^; [12]). Degradation of SPD-5 resulted in an even greater reduction of intestinal cell nuclei (Figure 1E and 1F, SPD-5^gut(-)^), perhaps due to a more direct effect on spindle formation. Surprisingly, degradation of SPD-2 or PCMD-1 had no impact on intestinal nuclear number, suggesting no perturbation in spindle formation or centriole duplication (Figure 1E and 1F, SPD-2^gut(-)^ and PCMD-1^gut(-)^). In contrast to their requirement in the *C. elegans* zygote, these results indicate that SPD-2 and PCMD-1 are not essential for mitotic centrosome function in embryonic divisions.

### SPD-5 is maintained at the ciliary base of C. elegans cilia

Specific roles for PCM proteins or subcomplexes during mitosis are difficult to distill as the PCM is rapidly evolving and under heavy regulation [18]. Thus, we sought to find interphase PCM that retains MTOC function to probe the role of distinct PCM proteins and/or subcomplexes. In many differentiated cells types, MTOC function is reassigned to a non-centrosomal site as part of the normal process of differentiation [19]. Cells with cilia provide an exception to this general phenomenon as the basal body derived from the centrosome often retains MTOC activity [20,21]. In *C. elegans*, only a subset of sensory neurons are ciliated and previous work found that after migration of the centriole to the cell surface to template the growth of the cilium, the centriole is degraded beginning with the loss of SPD-2 and followed by the removal of centriole cartwheel components such as SAS-4/CPAP [1,2,22]. As we would have expected the PCM to be removed with the centriole, we were therefore surprised to find endogenous SPD-5 localized to the ciliary base (CB) of all ciliated sensory neurons in adult worms alongside GIP-1 (Figure 2A). Analysis of SPD-5 localization during embryonic ciliogenesis in the amphid neurons together with SPD-2, SAS-4, and microtubules revealed that SPD-5 is maintained at the CB through the entire process of ciliogenesis despite the previously described [2] loss of SPD-2 and SAS-4 (Figure 2B and 2C). This pattern of localization suggests that the CB in *C. elegans* is composed of centriole-less PCM.

**Figure 2.**
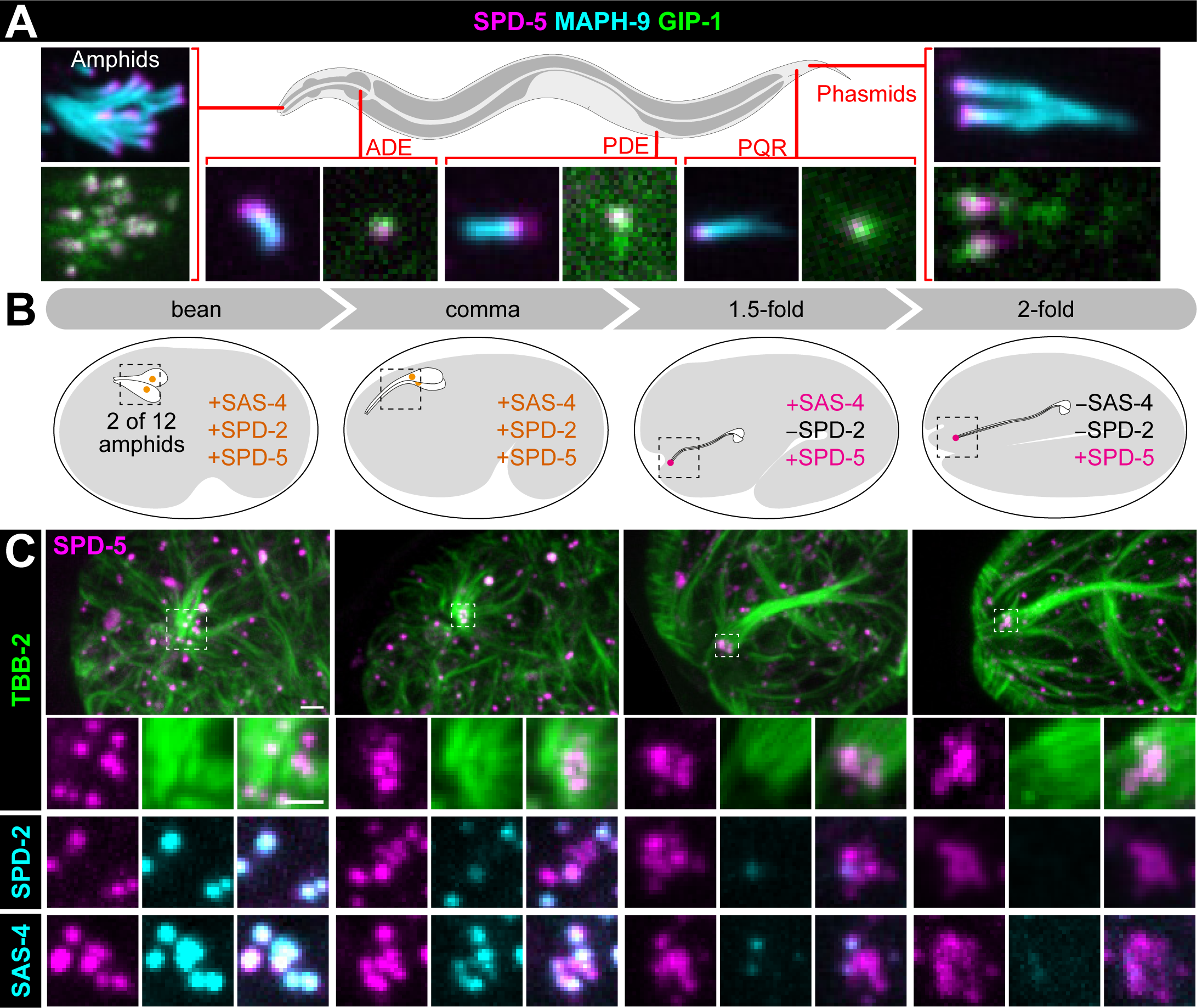
SPD-5 is maintained at the ciliary base while other centriolar and PCM protein are lost A) Localization of SPD-5 (magenta) in indicated ciliated sensory neurons relative to the axoneme (MAPH-9, cyan) and ciliary base (GIP-1, green) in *C. elegans* adults. B) Cartoon depicting the stages of amphid ciliogenesis in the embryo. Centrioles (orange) migrate to the tip of the dendrite during bean and comma stage and subsequently lose (magenta) SPD-2 at 1.5-fold stage, followed by SAS-4 at 2-fold stage. C) Localization of endogenously tagged indicated proteins in live embryos. Magnified views of the cluster of centrioles in the amphid neurons (12 total) shown below. Scale bar= 2µm; 1µm for insets.

### The ciliary base presents a unique MTOC organization

The CB has previously been suggested to function as an MTOC; γ-tubulin localizes near the CB and EBP-2/EB3 comets have been shown in the dendrite moving away from the cilium toward the cell body, however the structure imparting MTOC function was unclear from previous analysis [10,23]. To determine if the centriole-less PCM at the CB acts as an MTOC, we characterized the CB structure and microtubule organization in the adult phasmid neurons, a more isolated pair of ciliated sensory neurons. Localization of endogenous TBB-2/β-tubulin, GIP-1 and EBP-2 suggested that dynamic microtubules emanate directly from the CB; GIP-1 colocalized with SPD-5 at the CB and EBP-2 comets appeared to originate directly from the SPD-5 containing region (Figure 3A and 3B). In contrast to their localization to mitotic PCM, AIR-1/Aurora A and PLK-1/Plk1 did not localize to the centriole-less PCM at the CB (Figure S2).

**Figure 3.**
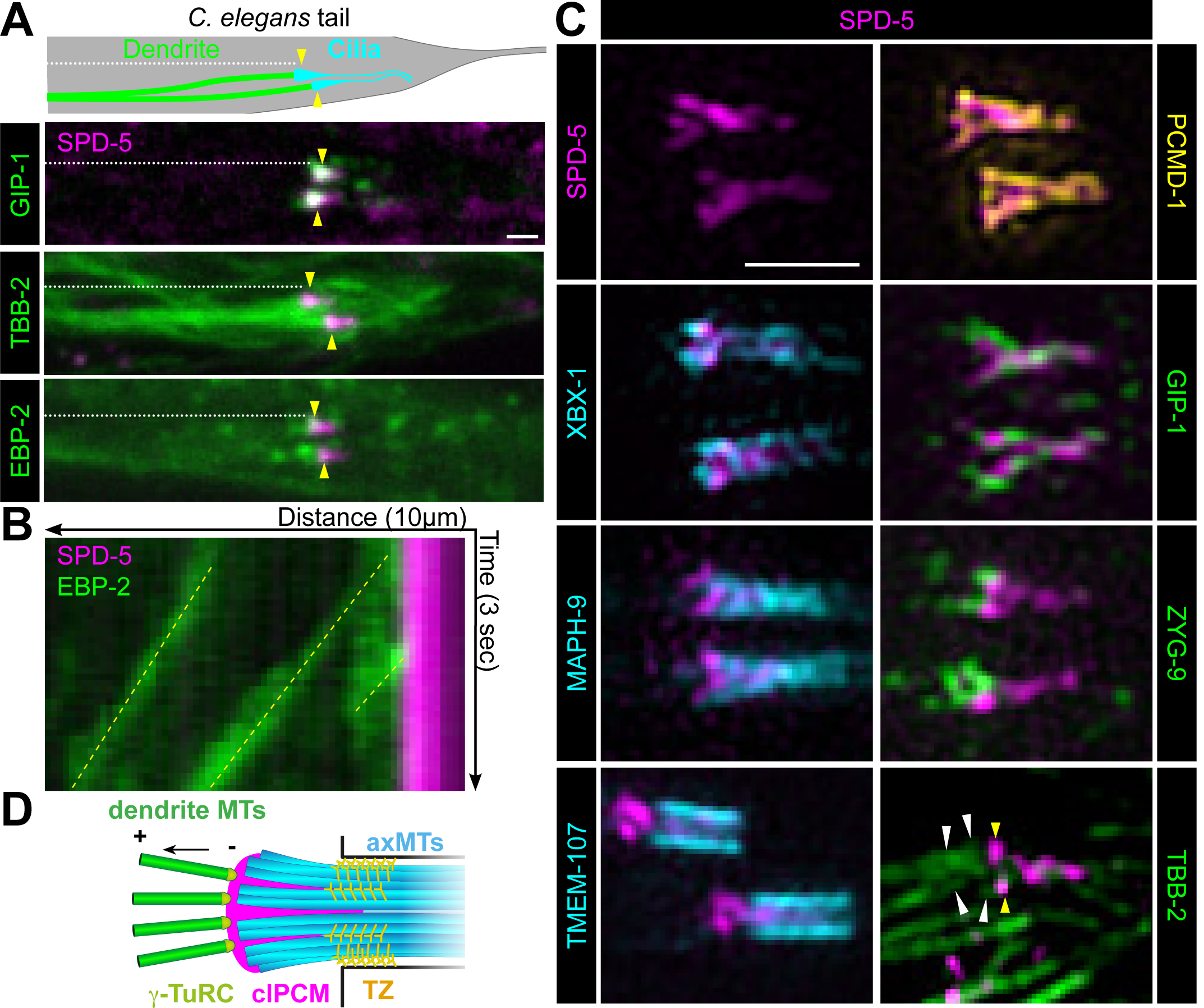
*Centriole-less PCM at the ciliary base serves as a MTOC*, see also Figure S2 A) Cartoon (top) and images from live adult worms (bottom) of the *C. elegans* phasmid neurons with ciliary base indicated (yellow arrowhead). B) Kymograph of EBP-2 comets (green, yellow dotted line) emerging from the ciliary base (SPD-5, magenta). C) 3D-SIM analysis of SPD-5 (magenta) localization at the ciliary base in phasmids cilia relative to PCMD-1 (yellow), axonemal (cyan), and microtubule-related (green) proteins; Yellow arrowheads indicate centriole-less PCM and white arrowheads indicate cytoplasmic microtubules. D) Cartoon summarizing the organization of the ciliary base as revealed by 3D-SIM. Scale bars= 1µm.

We further characterized the CB structure using 3D-SIM (Figure 3C), revealing that SPD-5 localizes in a ‘T’-shape, projecting inside the luminal space of the axoneme: SPD-5 localizes inside the barrel of the axoneme (XBX-1/ DNC2L1 and MAPH-9/MAP9) and transition zone (TMEM-107/TMEM107), together with PCMD-1, consistent with previous anecdotal reports of PCMD-1 localization in ciliated sensory neurons [7]. GIP-1 appeared to localize heterogeneously around and intermingled with the SPD-5 T-shape structure. The microtubule associated protein ZYG-9/chTOG was restricted to the dendritic space juxtaposed to SPD-5. Importantly, individual dendritic microtubules appeared to directly connect to the SPD-5 structure (Figure 3C, arrowheads, TBB-2). These observations revealed that microtubules are organized from a T-shaped centriole-less PCM and extend from the CB toward the cell body likely stabilized in the dendritic spaces by microtubule associated proteins such as ZYG-9. These data also suggest that the CB provides a minimal testbed to isolate and study roles of PCM proteins in MTOC function divorced from their other centrosomal functions.

### SPD-5 based centriole-less PCM drives MTOC function at the CB and ciliogenesis

To test whether PCM components localize interdependently at the CB and have a role in MTOC function, we degraded endogenous proteins specifically in ciliated sensory neurons following centriole migration and subsequent docking at the membrane at the onset of ciliogenesis and transition zone establishment (Figure 4A and S3, ‘csn(-)’). SPD-5 degradation caused a significant reduction of PCMD-1 and the complete loss of GIP-1 from the CB (Figure 4B and 4D), while PCMD-1 degradation resulted in the incomplete loss of SPD-5 and GIP-1 (Figure 4C and 4D). These data suggest that SPD-5 and PCMD-1 are present at the CB in different subcomplexes and that only a fraction of the centriole-less PCM is a complex of SPD-5 and PCMD-1.

**Figure 4.**
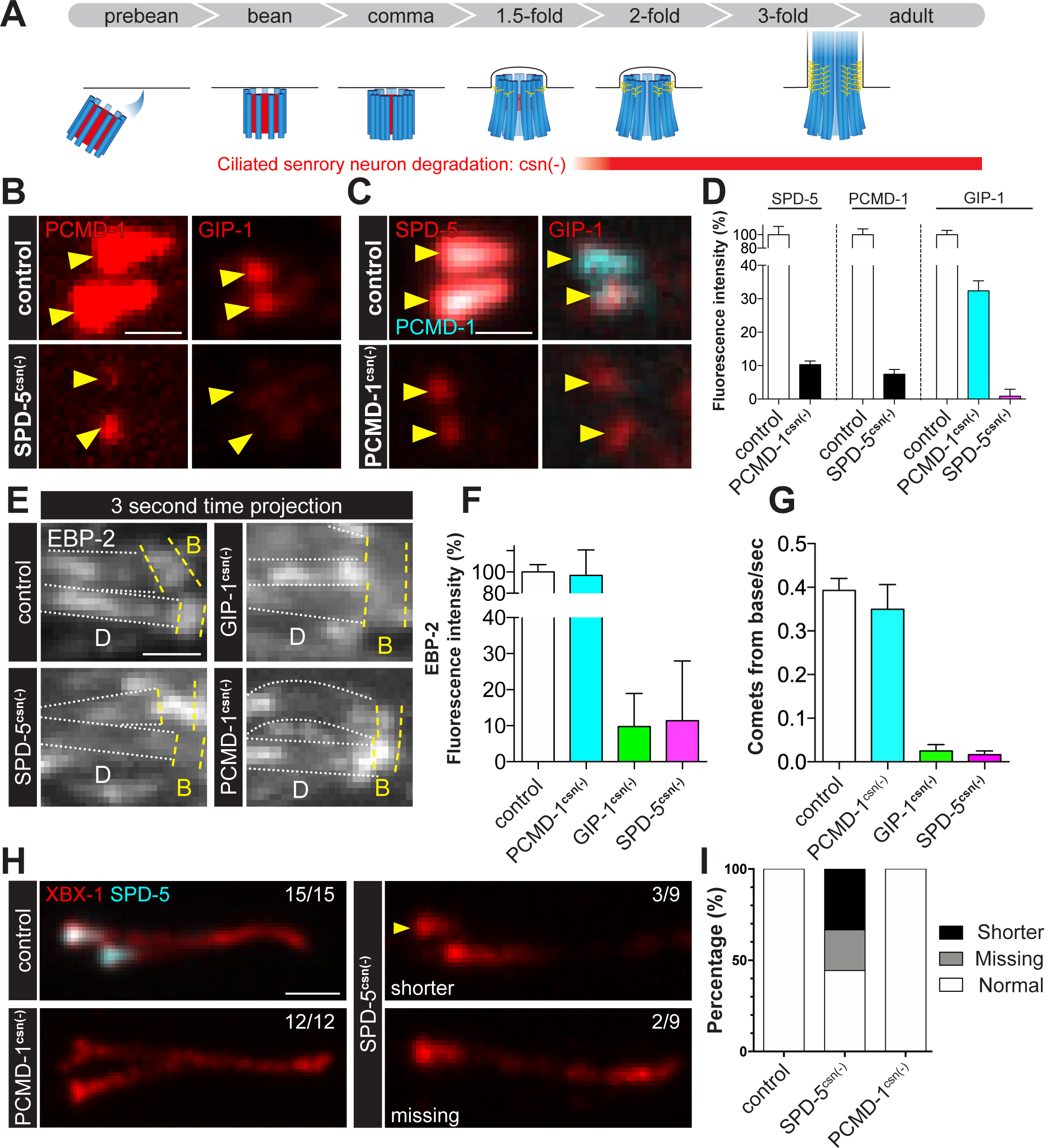
SPD-5 drives MTOC function at the ciliary base and ciliogenesis, see also Figure S3. A) Ciliogenesis steps based on previous reports during *C. elegans* development with centriolar microtubules (blue), internal centriole structures (red), and transition zone fibers (yellow) indicated. The *osm-6* promoter is used to drive ciliated sensory neurons specific degradation (csn(-)). B-D) Localization of indicated proteins at the ciliary base (yellow arrowhead) of phasmid cilia following SPD-5 or PCMD-1 depletion. D) Analysis of fluorescence intensity of indicated proteins following degradation of SPD-5 or PCMD-1. SPD-5: control= 100±12.45%, n= 10; PCMD-1^csn(-)^= 10.32±1.06%, n= 14; PCMD-1: control= 100±8.83%, n= 11; SPD-5^csn(-)^= 7.45±1.39%, n= 12; GIP-1: control= 100±6.41%, n= 10; PCMD-1^csn(-)^= 32.36±3.01%, n= 10; SPD-5^csn(-)^= 0.82±2.09%, n= 10. Values are mean ± standard error of mean. E) 3 second time-projection of EBP-2 comets along the dendrites (outlined by white dotted lines, ‘D’) near the ciliary base (yellow dotted lines, ‘B’) of phasmids cilia. F) EBP-2 fluorescence intensity relative to control: control= 100± 6.87%, n= 10; PCMD-1^csn(-)^= 96.85±23.8%, n= 10; GIP-1^csn(-)^= 9.74±9.25%, n= 10; SPD-5^csn(-)^= 11.35±16.6%, n= 10. G) EBP-2 comets emanating from ciliary base: control= 0.40±0.03 comets/sec., n= 10; PCMD-1^csn(-)^= 0.35±0.06 comets/sec., n= 10; GIP-1^csn(-)^= 0.03±0.01 comets/sec., n= 10; SPD-5^csn(-)^= 0.02±0.01 comets/sec., n= 10. H) Dynein protein XBX-1 localization (red) in indicated genotypes. Shorter cilium indicated by yellow arrowhead. I) Quantification of observed structural defects following SPD-5 or PCMD-1 degradation. Control: 100% normal; SPD-5^csn(-)^: 44.4% normal, 22.2% missing, and 33.3% shorter; PCMD-1^csn(-)^: 100% normal.

The presence of distinct subcomplexes of protein at the CB suggested that, like at the centrosome, these subcomplexes could also be functionally distinct. Furthermore, the partial loss of SPD-5 and γ-TuRC following PCMD-1 degradation suggested that PCMD-1 localizes a specific subcomplex of SPD-5 that is not required for MTOC function. To test the role of these subcomplexes in MTOC function at the CB, we monitored the movement of EBP-2 comets from the CB following degradation. In control, we observed an enrichment of EBP-2 at the CB with comets projecting from the CB toward the cell body (Figure 4E, G; Video S1). Upon degradation of SPD-5 or GIP-1, we observed a loss of both EBP-2 enrichment and comets, while PCMD-1 degradation did not affect either (Figure 4E and 4G; Video S1). Together, these results indicate that γ-TuRC is recruited by a SPD-5 dependent subcomplex of the centriole-less PCM to drive MTOC function at the CB.

Finally, we tested whether centriole-less PCM is required for ciliogenesis. SPD-5, but not PCMD-1, degradation led to variable defects in ciliary structure ranging from shorter to absent cilia, as visualized by an endogenously tagged dynein component, XBX-1 (Figure 4H and 4I). The phenotypic variability is likely due to variation in the timing of degradation but is consistent with a role for SPD-5 and the centriole-less PCM in establishing and/or maintaining the ciliary axoneme (Figure 4H and 4I).

## Discussion

Here, using multiple different cellular contexts, we found evidence for a SPD-5-based module that controls MTOC function within the PCM. This claim is supported by three lines of evidence. First, SPD-5 remains associated with dynamic microtubules and microtubule associated proteins in sub-PCM fragments that are spatially distinct from other PCM proteins such as SPD-2 and PCMD-1. Second, SPD-5, but not SPD-2 or PCMD-1, is essential for mitosis throughout development. Third, SPD-5 remains associated at the base of cilia with centriole-less PCM, a structure we argue represents the “pure” MTOC function of the centrosome. As loss of both γ-TuRC and SPD-5 from ciliated sensory neurons inhibited microtubule growth from the CB, we propose that SPD-5 alone recruits γ-TuRC to build and localize microtubules. As the SPD-5 functional homologs Centrosomin and CDK5RAP2 can directly interact with γ-TuRC, SPD-5 might serve as a platform to directly recruit γ-TuRC. Extensive studies have shown that the ability of the centrosome to expand its PCM is controlled by phosphorylation by mitotic kinases [24]. However, we did not observe AIR-1 or PLK-1 at the CB, indicative of the ability of SPD-5 to recruit microtubule regulators independent of this canonical mitotic regulation and making the CB a unique platform to study MTOC regulation.

The fact that SPD-2 and PCMD-1 are dispensable for embryonic cell divisions is in sharp contrast to their essential function in the first cell division of the *C. elegans* zygote. Although surprising, our work is in agreement with reports of phenotypes observed for *spd-2* [25] and *pcmd-1* temperature sensitive mutants [7], indicating that the essential nature of SPD-2 and PCMD-1 is due to their specific function in the zygote. We speculate that this zygote-specific requirement is due to the unique nature of the first mitosis of the embryo where the heavily repressed sperm centrosome must be reactivated in order to build the spindle [26]. Thus, SPD-2 and PCMD-1 may act as a zygote-specific catalyst for PCM assembly as has been suggested from in vitro studies [27]. Although the centrosome in other embryonic divisions in inactive as an MTOC in interphase, especially once embryonic cell cycles achieve gap phases [28,29], a residual shell of PCM in interphase could serve the same purpose [4,6,30].

Despite the fact that ultrastructural and localization studies found that the centrioles and SPD-2 are lost from the CB, we found that SPD-5 and PCMD-1 are maintained at the base of *C. elegans* cilia. Given that SPD-5, but not SPD-2, is stripped from the centrosome during mitotic exit into packets, the centriole-less PCM at the CB could be a stabilized remnant of PCM packets. However, we found that PCMD-1 does not localize to packets, suggesting the involvement of other mechanisms to maintain specific proteins at the CB. These remaining proteins could be protected from removal by dephosphorylation by mitotic phosphatases [5] or by proteasome-dependent degradation. Alternatively, or perhaps in addition, SPD-5 and PCMD-1 could be maintained through continued expression. Interestingly, SPD-5, but not PCMD-1 or SPD-2, contain an X-box motif in their promoter region, which is a conserved signature of genes regulated by ciliogenic transcription factors and indicating continued expression of *spd-5* specifically in ciliated sensory neurons [31]. We hypothesize that the combination of targeted protein degradation and transcriptional regulation, drive the specific loss or maintenance of centrosomal proteins. Altogether, we described a novel organization of PCM proteins, independent of the centriole, where SPD-5 drives MTOC function. These studies reveal the innate ability of SPD-5 to recruit microtubule regulators which likely underlies the ability of the centrosome to function as an MTOC.

## Materials and Methods

### C. elegans strains and maintenance

*C. elegans* strains were maintained at 20°C unless otherwise specified and cultured as previously described [32]. Strains used in this study are as follows

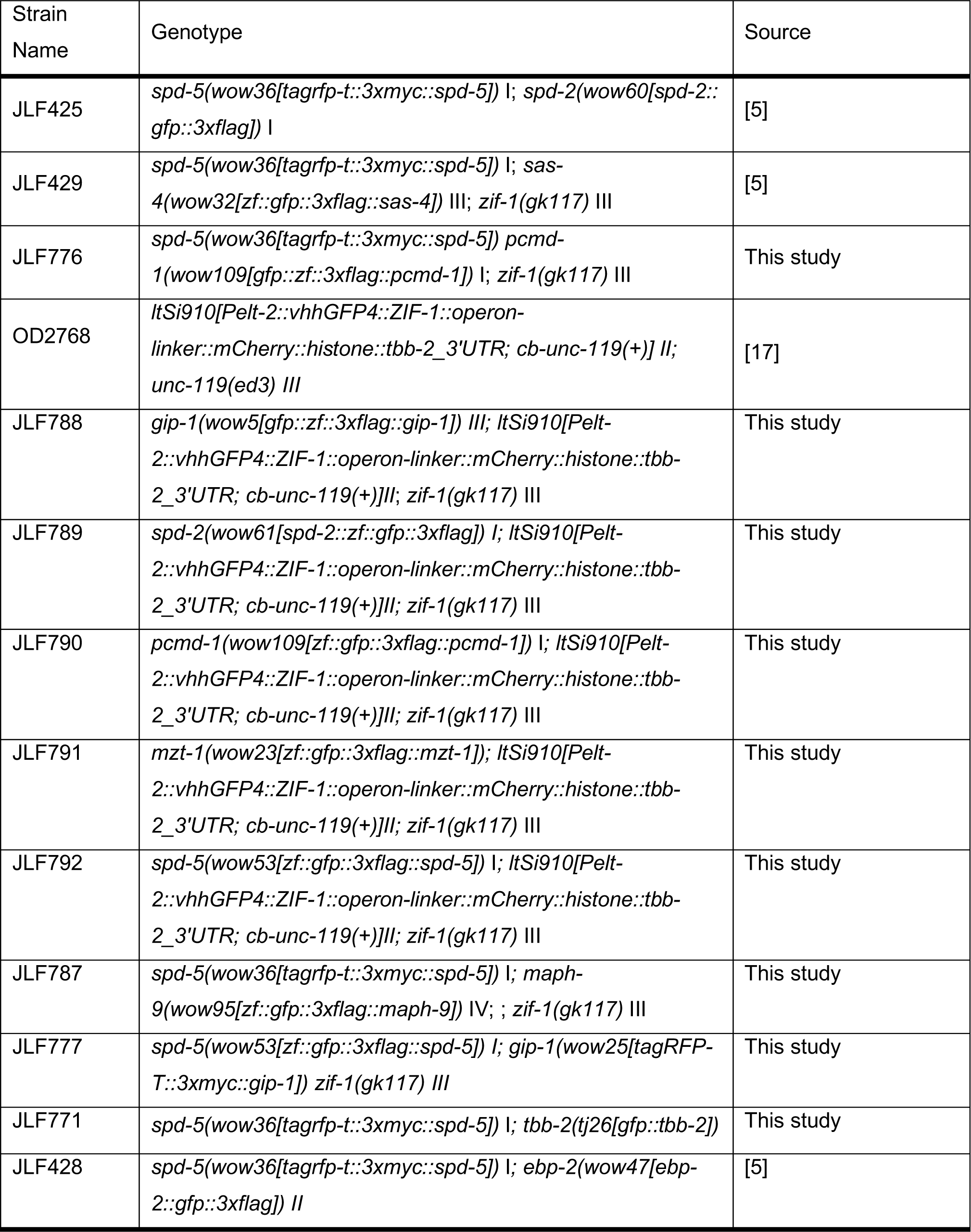
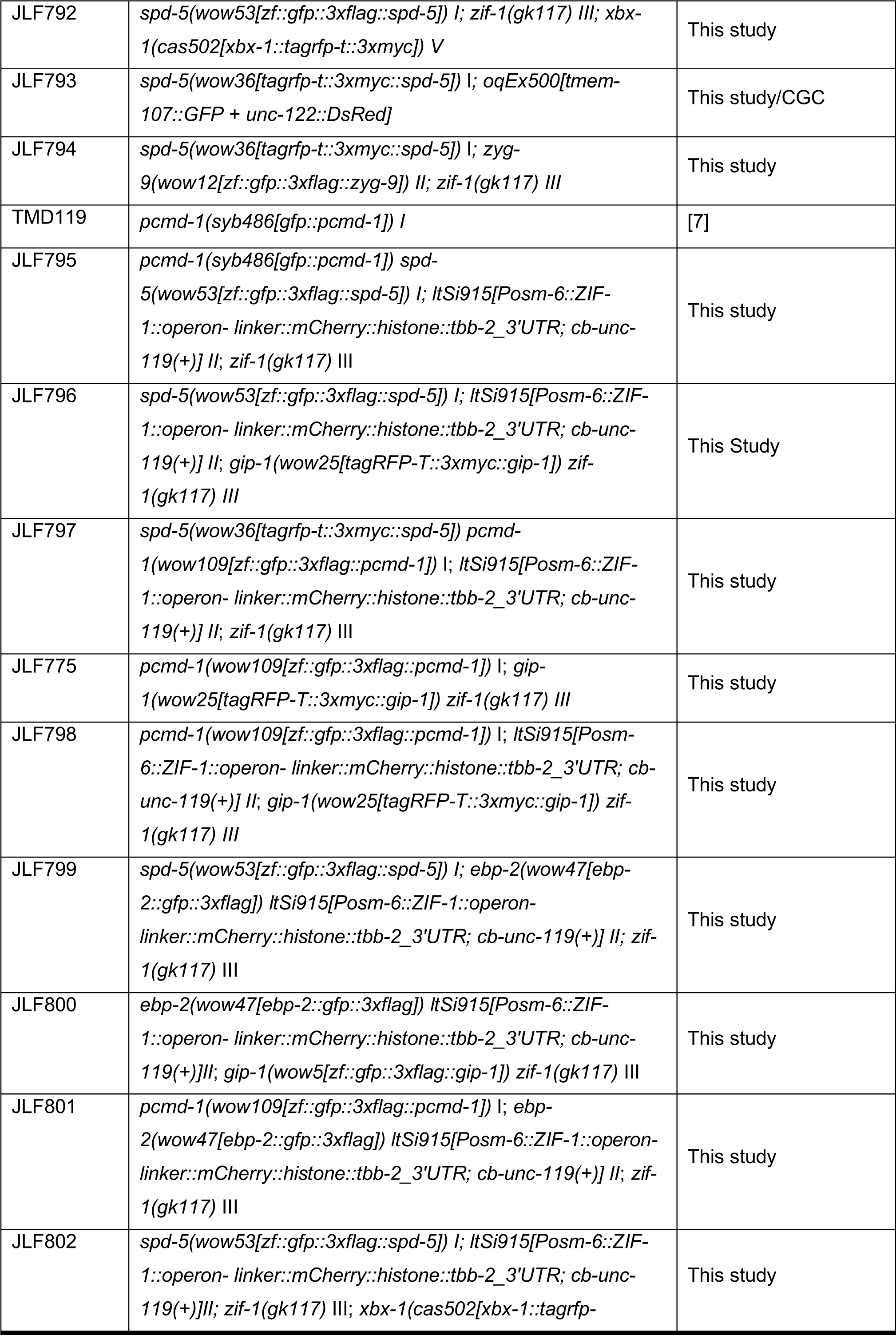
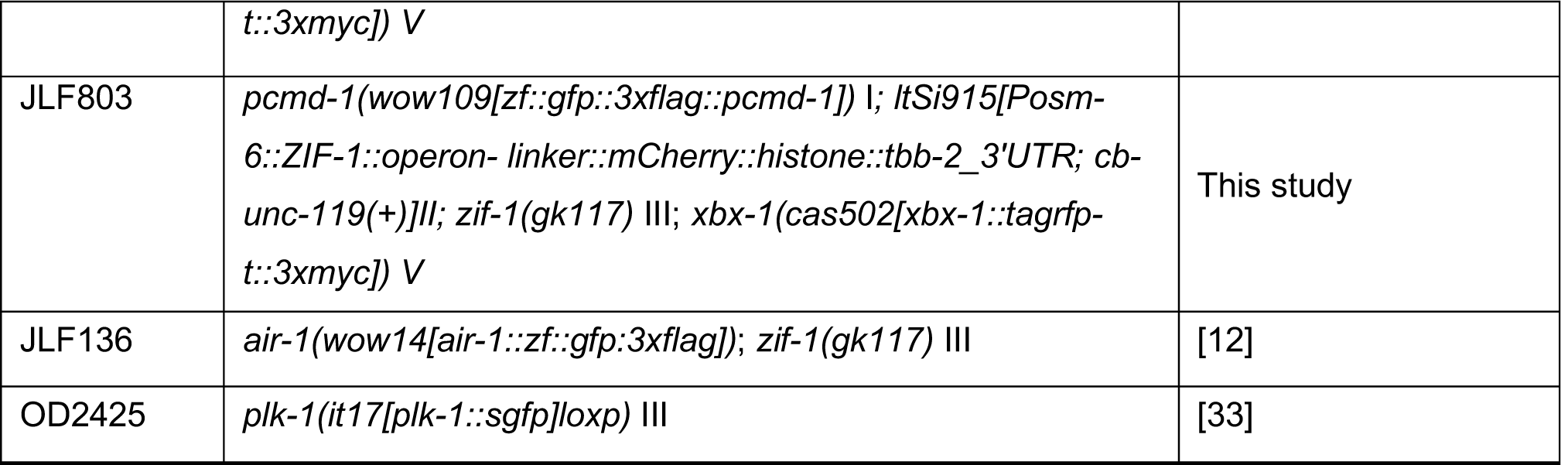

### CRISPR/Cas9

Endogenously tagged proteins used in this study were generated using the CRISPR Self Excising Cassette (SEC) method that has been previously described [34]. DNA mixtures (sgRNA and Cas9 containing plasmid and repair template) were injected into young adults, and CRISPR edited worms were selected by treatment with hygromycin followed by visual inspection for appropriate expression and localization [34]. sgRNA and homology arm sequences used to generate lines are as follows:

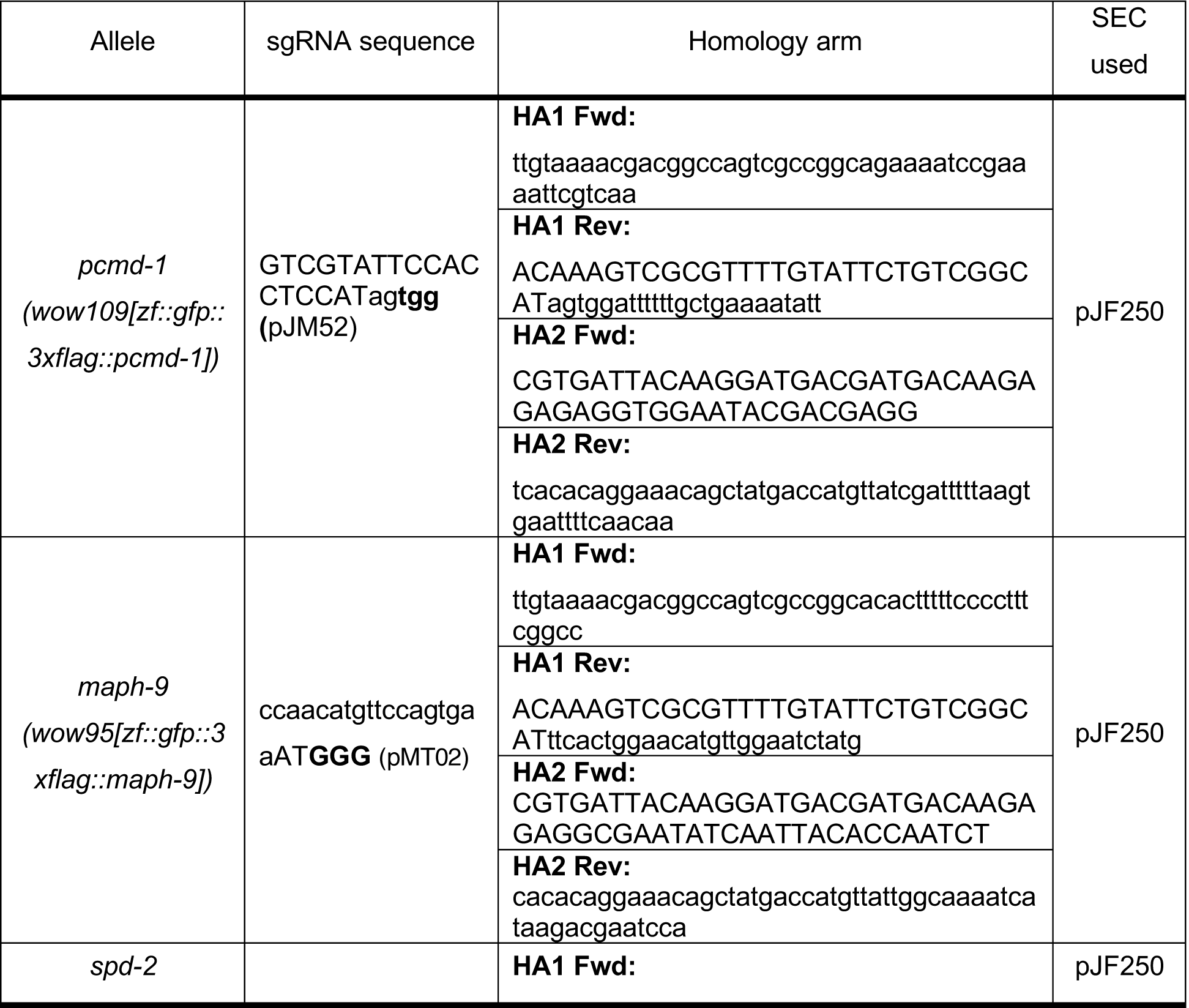
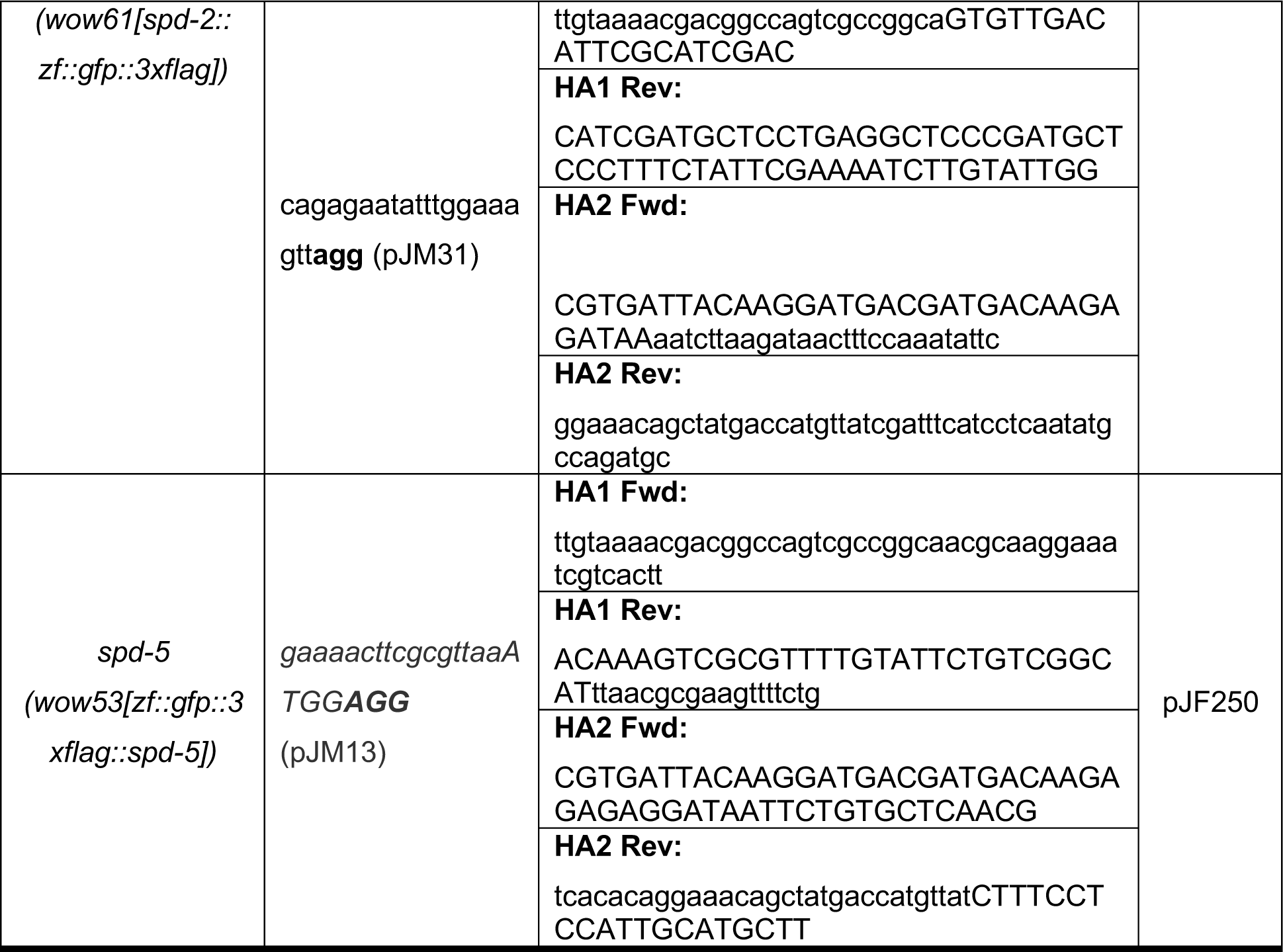

### Image acquisition

Worms were mounted on a pad (5% agarose dissolved in M9) immersed in 1mM Levamisol sandwiched between a microscope slide and no. 1.5 coverslip. The same technique was applied without Levamisole for mounting embryos. Spinning-disk confocal images were acquired on a Nikon Ti-E inverted microscope (Nikon Instruments) equipped with a 1.5x magnifying lens, a Yokogawa X1 confocal spinning disk head, and an Andor Ixon Ultra back thinned EM-CCD camera (Andor), all controlled by NIS Elements software (Nikon). Images were obtained using a 60x (NA= 1.4) or 100x Oil Plan Apochromat objective (NA= 1.45). Z-stacks were acquired using a 0.2 µm step. For EBP-2 comet imaging, images were acquired as a stream of single plane images focused on the ciliary base, with a 100ms exposure. Structured illumination microscopy images were acquired on an OMX BLAZE V4 microscope (GE) belonging to Stanford Cell Science Imaging Facility equipped with an U-PLANAPO 100x (NA= 1.4). objective and Photometrics Evolve 512 EM-CCD cameras, all controlled by DeltaVision software. Structured Illumination images were generated using SoftWoRx software. Images were adjusted for brightness and contrast using ImageJ software.

#### Image Quantification

### PCM intensity measurement

Fluorescence intensity was measured by defining a square image stack 20 pixels wide x 11 images deep around the phasmids. Another stack of the exact same dimensions was generated in the neighboring tissue. Both stacks were sum projected and the phasmid intensity was measured by subtracting the total intensity of the neighboring sum projection from the total intensity of the phasmids sum projection.

### EBP-2 intensity measurement

Fluorescence intensity was measured by defining a round Region of Interest of 10 pixels in diameter around the phasmid in the single plane. Another ROI of the exact same dimensions was generated in the neighboring tissue. The base of cilia intensity was measured by subtracting the intensity of the neighboring tissue from the intensity of the base of cilia.

### Statistics

Statistical analyses were performed using R and Prism (GraphPad software, La Jolla, Ca, USA).

## Acknowledgements

We thank Karen Oegema, Tamara Mikeladze-Dvali, Dan Dickinson, and Bob Goldstein for providing strains and CRISPR advice and protocols. We also thank Kang Shen, Piali Sengupta, and members of the Feldman lab for helpful discussions about the manuscript. Some of the nematode strains used in this work were provided by the *Caenorhabditis* Genetic Center, which is funded by the NIH Office of Research Infrastructure Programs (P40 OD010440). This work was supported by an NIH New Innovator Award DP2GM119136-01 and R01GM136902 awarded to J.L.F. J.M. is supported by an American Heart Postdoctoral Fellowship. The project described was also supported, in part, by Award Number 1S10OD01227601 from the National Center for Research Resources (NCRR). Its contents are solely the responsibility of the authors and do not necessarily represent the official views of the NCRR or the National Institutes of Health.

## Author Contributions

Conceptualization, J.L.F and J.M.; Methodology, J.M., J.L.F., S.E. and M.V.T.; Formal Analysis J.M.; Investigation, J.M., J.L.F., S.E. and M.V.T.; Writing -Original Draft, J.L.F and J.M.; Writing – Review & Editing, J.L.F and J.M.; Visualization, J.L.F and J.M.; Supervision, J.L.F.; Funding Acquisition, J.L.F and J.M.

## Supplemental Figure Legends

**Figure S1.**
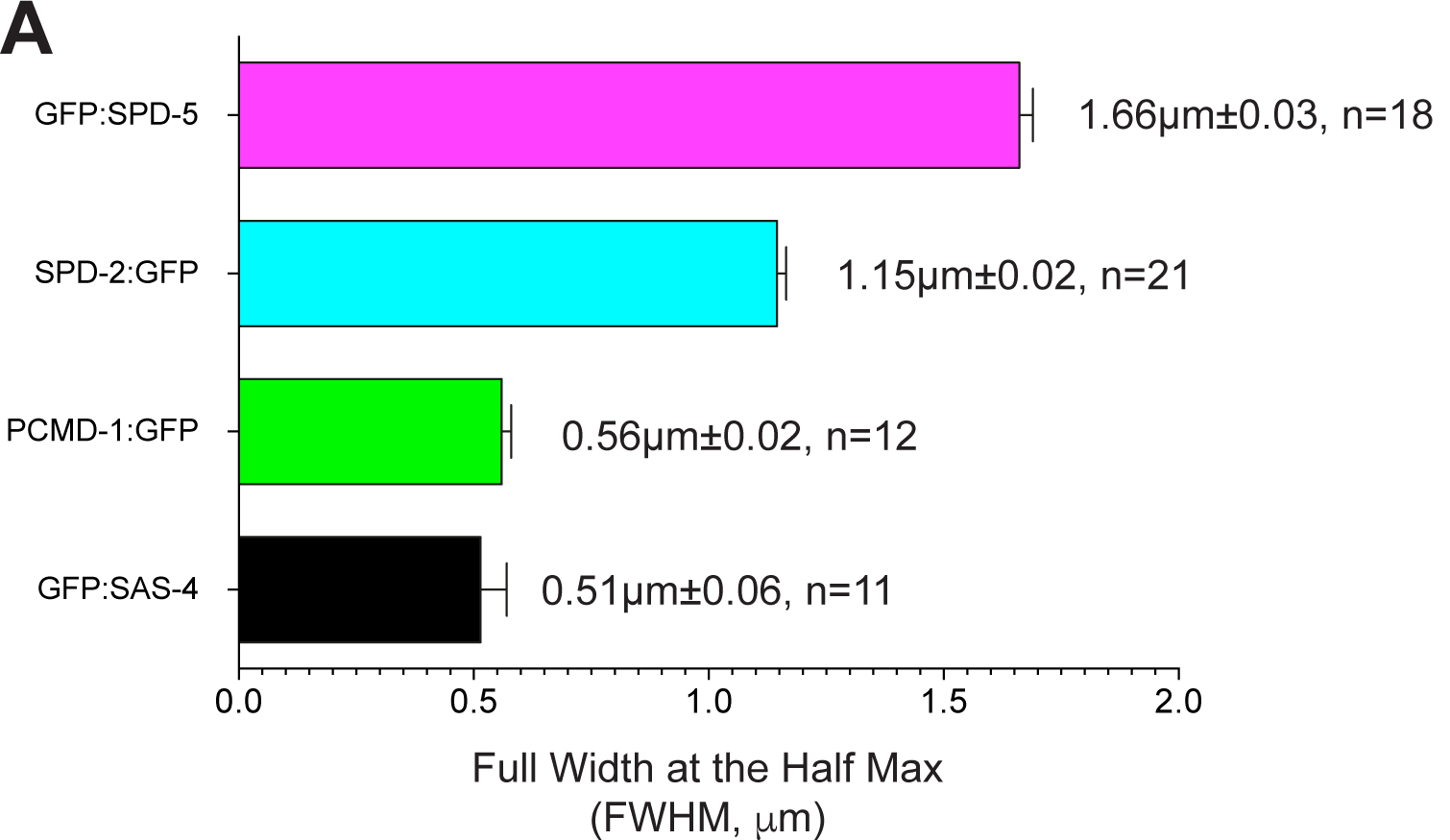
PCMD-1 localizes in between the centriolar protein SAS-4 and PCM protein SPD-2, related to Figure 1. A) Average width of pixel intensity profile for each protein at ABp centrosome at nuclear envelope breakdown: SPD-5: 1.66±0.03µm, n= 18; SPD-2: 1.15±0.02µm, n= 21; PCMD-1: 0.56±0.02µm, n= 12; SAS-4: 0.51±0.06µm, n= 19.

**Figure S2.**
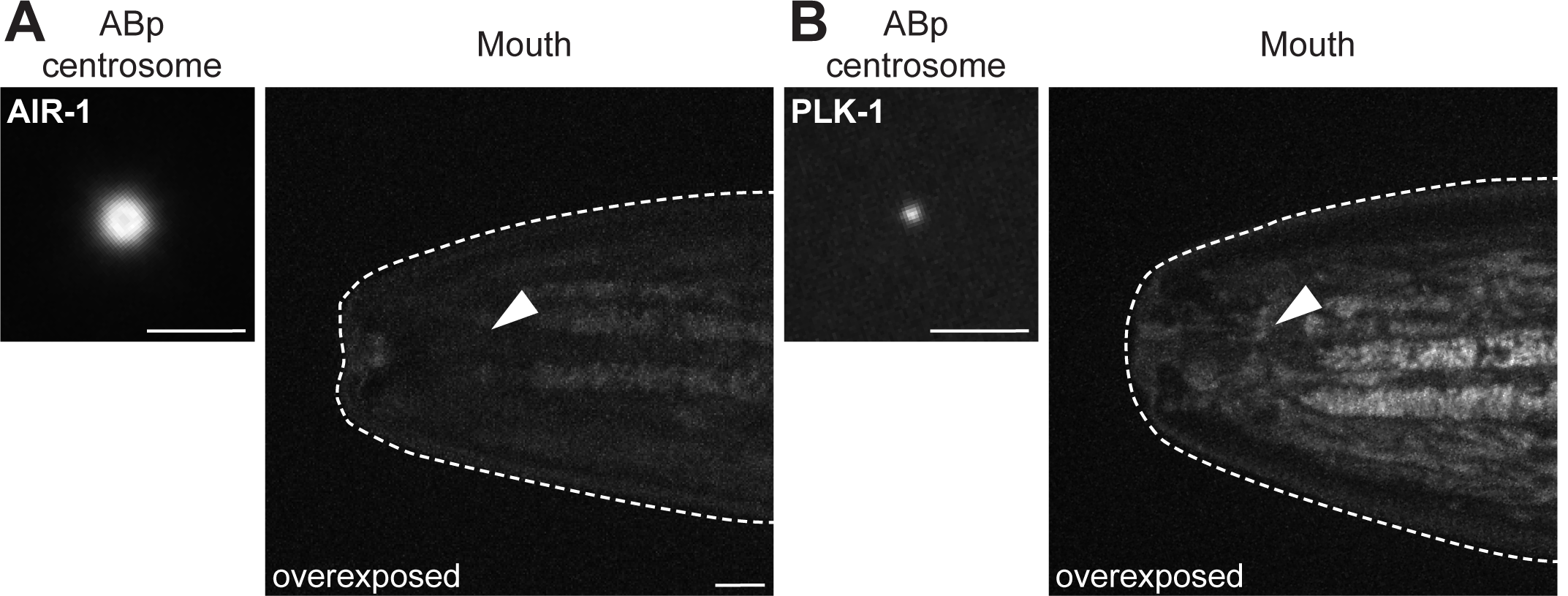
AIR-1 and PLK-1 kinases are absent at the ciliary base of ciliated sensory neurons, related to Figure 3. A-B) Live confocal imaging of indicated proteins in the ABp centrosome from a 4-cell embryo at nuclear envelope breakdown (NEBD) or in the adult mouth (delimited by the dashed line). Scale bar = 5µm.

**Figure S3.**
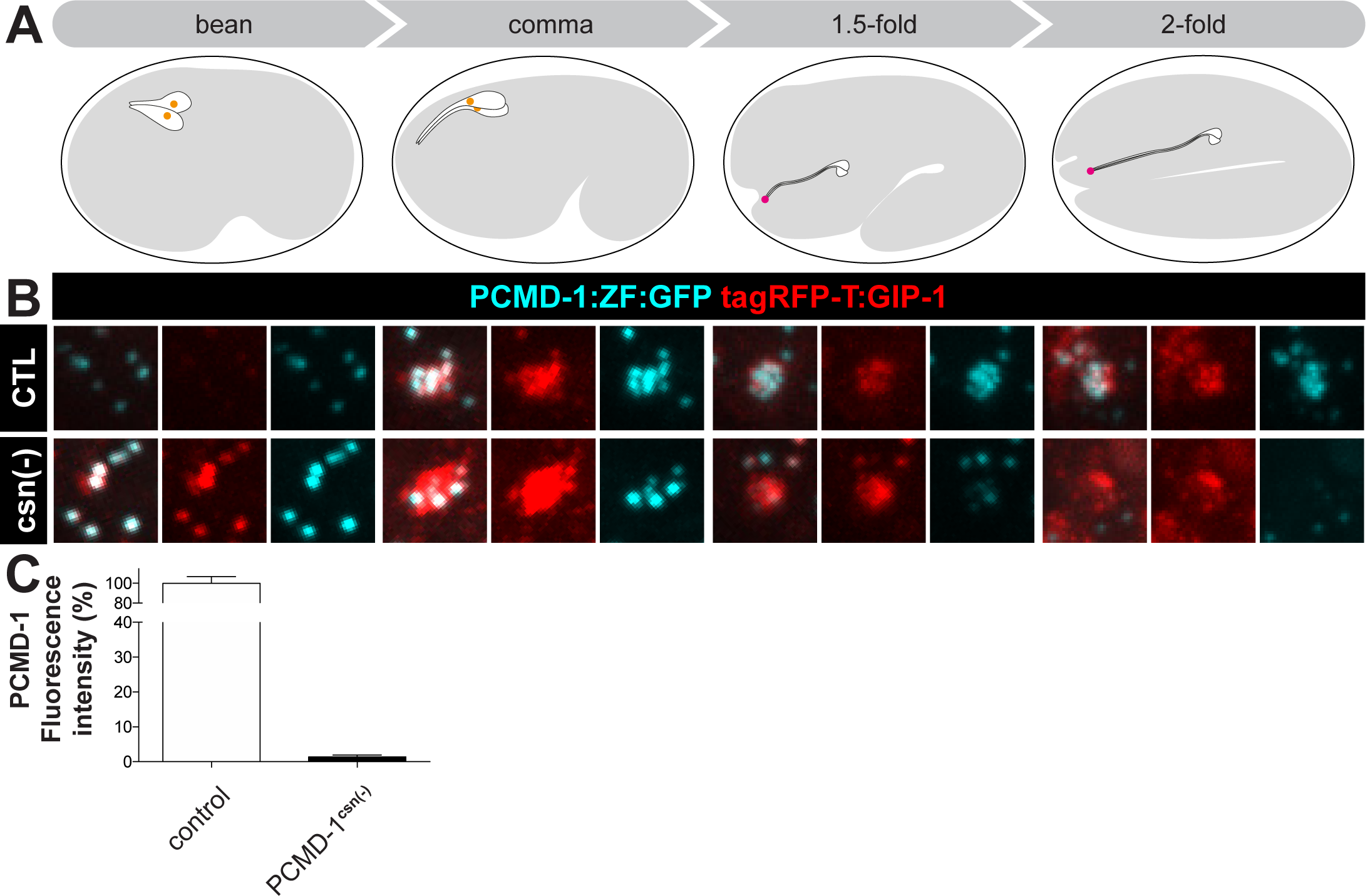
*osm-6* promoter drives ciliated sensory neuron specific degradation of proteins at the onset of ciliogenesis, related to Figure 4. A) Cartoon depicting the different stages of *C. elegans* ciliogenesis in amphid neurons from bean (left) to 2-fold stage (right). B) Analysis of PCMD -1 upon ciliated sensory neurons specific degradation (bottom row) compared to a control (top row). GFP signal reduction begins at the 1.5-fold stage and is completely gone at 2-fold. C) Analysis of fluorescence intensity of PCMD-1 following degradation in adult phasmids. control= 100±6.85%, n= 10; PCMD-1^csn(-)^= 1.40±0.53%, n= 10. Values are mean ± standard error of mean.

Video S1. EBP-2 comets upon ciliated sensory neuron specific degradation of PCM proteins, related to Figure 4.

Yellow circle represents the position of the ciliary base. Contour of the phasmids are delimited by the yellow dashed line.

